# Ventral tegmental area dopamine controls timing variability

**DOI:** 10.1101/2025.10.27.684913

**Authors:** Matthew A. Weber, Kartik Sivakumar, Alexandra S. Bova, Ervina E. Tabakovic, Mackenzie M. Conlon, Mayu Oya, Rachel C. Cole, Arturo I. Espinoza, Youngcho Kim, Nandakumar S. Narayanan

**Author notes:** **Corresponding Author** Nandakumar Narayanan.

## Abstract

Precision defines successful behavior, yet the brain mechanisms promoting precision are unclear. Here, we dissect dopaminergic circuits controlling precision along a single behavioral dimension – the timing of action. We use an interval timing paradigm that requires participants to indicate their estimate of an interval of several seconds with a motor response. We find that humans with Parkinson’s disease (PD) had increased timing variability that predicted PD-related cognitive deficits and executive dysfunction. Surprisingly, lesioning ventral tegmental area (VTA) dopamine neurons increased temporal variability. Further, GCaMP6s fiber photometry demonstrated that VTA dopamine neuron activity is strongly modulated at the start of temporal intervals, and that this trial start-related activity predicted temporal variability. Finally, we found that stimulation of VTA dopamine neurons improved timing by decreasing temporal variability in both intact and dopamine-depleted animals. Our data establish a model of cognitive symptoms of human PD and provide insight into the neuronal control of temporal variability, which impacts a wide range of executive functions.

## INTRODUCTION

To be effective, behavior must be accurate as well as precise. While much of behavioral neuroscience has focused on accurate behavioral outcomes, considerably less effort has been focused on behavioral precision and variability^1,2^. A neurotransmitter system that is critical for behavioral variability is dopamine, as drugs and diseases of dopamine can modulate behavioral variability^3–5^. However, it is unclear how dopamine modulates precise behavior, which is important to understanding a wide range of mammalian behavior as well developing new biomarkers and treatments for human diseases such as schizophrenia, attention-deficit hyperactivity disorders, substance use disorder, and Parkinson’s disease (PD)^6–8^.

Complexity can challenge attempts to quantify behavioral variability. We addressed this by turning to a paradigm configured to capture behavioral variability – interval timing^9^. This paradigm requires participants to estimate an interval of several second by making a motor response and measures temporal precision by quantifying the variability of responses^10^. Past work has shown that humans with disrupted dopamine, including schizophrenia and Parkinson’s disease, have increasing timing variability^11–16^, which implicates dopaminergic circuits in temporal variability.

We systematically investigated dopaminergic circuits and temporal variability. We found that PD patients had increased timing variability that predicted executive dysfunction and cognitive status. In mice, we found that depleting dopamine in the ventral tegmental area increased timing variability and that ventral tegmental area dopaminergic neuron activity triggered at the start of timing trials predicted timing variability. Further, we found that stimulating ventral tegmental dopamine neurons improved timing by decreasing timing variability in intact and dopamine-depleted animals. We interpret our data in the context of ventral tegmental dopamine and cognitive variability.

## METHODS

### Humans

We recruited 94 Parkinson’s disease (PD) patients and 43 health comparison control participants from the movement-disorders clinic at the University of Iowa according to methods described in detail previously^12,17^. Patients were examined by a movement-disorders physician to verify that they met the diagnostic criteria recommended by the United Kingdom PD Society Brain Bank. We obtained written informed consent from each participant (IRB Protocol 201707828). PD patients participated in the study while taking their prescribed medications, including levodopa, as usual. All testing was done in a laboratory setting in the University of Iowa’s Department of Neurology. All stimuli were presented via Psychtoolbox v3.0, a MATLAB toolbox running on a Dell XPS workstation and MATLAB 2018b, and a 19-inch monitor. We performed a peak interval timing task with 3-second and 7-second trials randomly intermixed^17^ that required participants to estimate a duration of either 3 seconds or 7 seconds after reading instructions displayed as text at the center of a video screen. White instructional text on the center of a black screen was presented that read “Short interval” on 3-second interval trials and “Long interval” on 7-second interval trials. Interval durations were never communicated to the patient. After the instructions were displayed for 1 second, the start of the interval was indicated by an imperative “Go” cue: an image of a solid box in the center of the computer screen. The cue was displayed on the screen for the entire trial, which lasted 8–10 s for 3-second intervals and 18–20 s for 7-second intervals. Participants were instructed to estimate the duration of the intervals without counting. As a distraction to discourage counting, vowels appeared at random intervals in the center of the screen. Participants were instructed to press the keyboard spacebar at the start of when they judged the target interval to have elapsed. On 15% of the trials, participants were then given nonnumerical/nonverbal performance feedback on the screen indicating their response relative to the actual target interval; feedback was a horizontal green line indicating how close their keypress was to the target time. Because feedback was relatively infrequent, we did not analyze this event. The time between trials varied from 3–8 seconds. All participants performed 6 training trials prior to testing trials and verbalized understanding of the task. Testing sets consisted of 40 trials of each interval length, for a total of 80 trials. Trials were presented in a random order of 20 trials blocks; four blocks were presented. Participants took a self-paced break between each block and moved to the next block by pressing any key. We analyzed the mean keypress time to capture timing accuracy for PD patients and control participants. To analyze timing precision, we computed the coefficient of variance (CV) by dividing the standard deviation by the mean keypress time. Of note, data from some participants was included in prior work^12,17^.

### Mice

All experimental procedures were approved by the Institutional Animal Care and Use Committee (IACUC) at the University of Iowa and performed in accordance with relevant guidelines of Protocol #3052039. Two different strains of mice were used: 1) wild type male and female C57BL/6 mice received from Jackson Labs (Bar Harbor, ME) at approximately 12–14 weeks of age; and 2) heterozygous male and female DAT^IRES^-Cre (B6.SJL-Slc6a3tm1.1(cre)Bkmn/J) mice bred in house from breeding pairs received from Jackson Labs (Strain #:006660). Prior to behavioral training or surgical procedures, all mice were communally housed on a 12-hour light/dark cycle with *ad libitum* access to laboratory rodent chow and water. To facilitate operant behavioral training, mice were individually housed and maintained on a restricted diet with *ad libitum* access to water.

### Interval timing switch task

We used a mouse optimized interval timing task described in depth previously^18–25^. Mice were trained in standard sound attenuated operant chambers (MedAssociates, St. Albans, VT) with a reward hopper located between the two nosepoke response ports (left and right) on the front wall and another nosepoke on the back wall. Each 90-minute session was randomly organized into 50% “short” 6 second interval trials and 50% “long” 18 second interval trials. Short and long trials had identical audio and visual cues and are self-initiated by a response at the back nosepoke port. In a short trial, mice must respond as the designated nosepoke after 6 seconds to then receive a 20 mg sucrose reward (BioServ, Flemington, NJ). This short designated nosepoke is counterbalanced left vs. right across mice. When a short trial is initiated, mice begin by responding at the designated nosepoke; mice may respond multiple times, but the reward is only delivered for the first response after 6 seconds. In a long trial, mice must respond at the opposite nosepoke, which is designated for long trials in which mice receive a reward for the first response after 18 seconds. Since cues are identical for both short and long trials, the optimal strategy is for mice to start responding at the designated short trial nosepoke, but if enough time passes with no reward delivered, the mice switch to the designated long trial nosepoke. The switch response time is defined as the moment when the mouse leaves the short trial nosepoke and is a measure of the rodent’s internal estimate of time, as in other interval timing tasks^19,21^. Only the long trials in which mice make an appropriate switch are analyzed^18–20,22–25^.

### Surgical procedures

Surgical methods were identical to those described previously^19,22,25,26^. Mice were anesthetized with 4.0% isoflurane at 400 mL/min and maintained on 1.5%–3.0% isoflurane at 120 mL/min (SomnoSuite, Kent Scientific, Torrington, CT, USA). Craniotomies were drilled above the ventral tegmental area (AP +3.3 mm / ML ±1.1 mm). Viruses, neurotoxins, or vehicles were infused over 10 minutes (0.05 µL/min; Legato 130 Syringe Pump, kd Scientific, Holliston, MA), with a 5-minute wait period before removing the needle. For fiber photometry and optogenetic experiments, at least three stainless steel screws were attached to the skull to stabilize head cap assemblies, which were sealed with cyanoacrylate (“SloZap,” Pacer Technologies, Rancho Cucamonga, CA) and accelerated with “ZipKicker” (Pacer Technologies) and methyl methacrylate (AM Systems, Port Angeles, WA). Following all surgical procedures, mice were allowed at least 1 week to recover before starting food restriction and interval timing training.

For dopamine depletion experiments^22^, wild type C57BL/6J mice were first trained on the interval timing switch task described above for approximately 3–4 weeks before surgical procedures. On the day of surgery, 6 hydroxydopamine hydrobromide (6-OHDA; Millipore Sigma #162957, Darmstadt, Germany) and desipramine hydrochloride (Millipore Sigma #D3900) were freshly made and stored on ice away from light. 6-OHDA was prepared at 2 mg/ml in 0.03% ascorbic acid (AA) and desipramine at 4 mg/ml in 0.9% saline. Desipramine (25 mg/kg) was injected intraperitoneally (IP) prior to starting surgical procedures to protect norepinephrine terminals against 6-OHDA^27–29^. Mice were randomly assigned to either the vehicle group or the ventral tegmental area dopamine depletion group. The vehicle group received microinjections of 0.5 μl 0.03% AA bilaterally, and the dopamine depletion groups received equal volumes of 6-OHDA prepared in AA (DV -4.6 mm at 10° laterally).

For fiber photometry experiments, a Cre-dependent GCaMP6s virus (Addgene, Watertown, MA; Catalog #100845-AAV5; pAAV.Syn.Flex.GCaMP6s.WPRE.SV40) was unilaterally injected in the VTA (DV -4.6 mm at 10° laterally) of DAT^IRES^-Cre mice. A fiber optic cannula (Doric Lenses Inc, Quebec, Cananda; NA 0.48) was then implanted in the VTA (DV -4.5 mm at 10° laterally).

For optogenetic stimulation experiments, a Cre-dependent channelrhodopsin^30^ (ChR2; Addgene; Catalog #20297-AAV5; pAAV-EF1a-DIO-hChR2(H134R)-mCherry) was bilaterally injected in the VTA (DV-4.6 mm at 10° laterally) followed by implantation of bilateral fiber optics (Doric Lenses Inc; NA 0.22; DV -4.5 mm at 10° laterally). There were two separate cohorts of optogenetic stimulation mice. In one cohort, only Cre-dependent ChR2 was injected in the VTA. In the other cohort, 6-OHDA was injected first using the same procedures as the dopamine depletion experiments described above; desipramine was also injected to preserve norepinephrine terminals. After 6-OHDA VTA injections, Cre-dependent ChR2 was injected and fiber optics implanted as described above.

### Fiber Photometry

Mice were extensively trained in the interval timing task described above before being acclimatized to fiber photometry procedures. Mice were unilaterally tethered to an optical patchcord connected to a fiber photometry system (Doric Lenses, Inc; Québec, Cananda) with two excitation wavelengths (405-nm LED isosbestic signal modulation at 208.616 Hz and 470-nm LED Ca^2+^-dependent GCaMP6s signal modulated at 572.205 Hz). Isosbestic and GCaMP6s fluorescence wavelengths were measured via integrated LED mini cubes and photodetector (iLFMC4, Doric Lenses, Inc). Changes in GCaMP6s fluorescence were downsampled to 100 Hz, low-pass filtered at 5 Hz, detrended to compensate for exponential decay and photobleaching, motion corrected and normalized via the fitted isosbestic signal to calculate ΔF/F_0_, z scored across the entire session, and aligned to task events^26,31^.

### Optogenetics

Similar to fiber photometry recordings, mice were extensively trained prior to optogenetic stimulation procedures. Mice were tethered to a 473-nm laser diode (Doric Lenses, Inc) through a bilateral optical rotary joint and optical patchcords (both from Doric Lenses, Inc)^19,32^. Laser power output was adjusted prior to each session to 12 mW at the tips of the optical patch cords. Optogenetic stimulation was controlled with custom written Arduino protocols and delivered during the first 2 seconds of randomly selected trials controlled by MedAssociates. Separate sessions were run with laser stimulation delivered at either 4 Hz or 20 Hz. We focused on interval timing switch trials and compared mean switch response time and switch response time CV when no laser light was delivered vs. when laser light was delivered.

### Histology

Mice were sacrificed following the completion of all experiments. Mice were deeply anesthetized using ketamine (100 mg/kg IP) and xylazine (10 mg/kg IP) and transcardially perfused with phosphate buffered saline (PBS) and 4% paraformaldehyde (PFA). Brains were removed and fixed in 4% PFA overnight, followed by immersion in 30% sucrose for approximately 48 hours, and then frozen in cryoprotectant at -80 °C. Brains were sliced at 40 µm coronal sections of the midbrain using a cryostat (Leica Biosystems, Deer Park, IL). Selected sections were then stained to visualize tyrosine hydroxylase, according to methods described previously^20,23^, as well as native fluorescence of GCaMP6s^26^ or ChR2^33,34^. All images were captured using VS ASW S6 imaging software (Olympus, Center Vallet, PA) to confirm histological locations of 6-OHDA dopamine depletion, GCaMP6s and ChR2 fluorescence, and fiber optic locations.

### Statistics

All data were analyzed with custom code written in MATLAB; all code and data will be available upon final publication at: http://narayanan.lab.uiowa.edu/article/datasets. The two primary measures of interval timing were quantified by 1) the mean switch response time of all switch trials performed during an interval timing session and 2) the switch response time coefficient of variance (CV) of all switch trials performed during an interval timing session. CV was calculated as the standard deviation divided by the mean and is expressed as a percentage. For VTA dopamine depletion experiment without optogenetic stimulation, post-surgery mean switch times and switch time CV were normalized to each mouse’s pre-surgical baseline to account for mouse specific variability in timing behavior. Differences between interval timing performance in either human or mouse data were computed using ANOVA from linear models (*fitglme* and *anova* in MATLAB). We accounted for human participant-specific or animal-specific variances in all analyses and set statistical significance at alpha 0.05. Pearson’s partial correlation (*partialcorr* in MATLAB) was used to determine the strength of relationships between human interval timing and both MOCA and Executive Function, controlling for UPDRS scores. Pearson’s partial correlation was also used to determine the strength of relationships between GCaMP6s ΔF/F_0_ and both mean switch response and switch response CV, controlling for multiple behavioral sessions from each mouse.

## RESULTS

We analyzed interval timing performance in 94 PD patients and 43 demographically-matched control participants in a two-interval timing task with 3- and 7-second intervals (Fig. 1A)^12^. The cumulative distribution of response times for each trial type (dashed lines: 3-second interval; solid lines: 7-second interval) are shown in Fig. 1B. We found that PD patients had greater mean response times in the 3-second interval timing task (Control: 3.61 ± 0.18 seconds vs. PD: 4.61 ± 0.23 seconds, *p* = 0.005; Fig. 1C) but not in the 7-second interval timing task (Control: 6.15 ± 0.15 seconds vs. PD: 6.41 ± 0.15 seconds, *p* = 0.267; Fig. 1C). However, PD patients had reliably increased coefficients of variance (CV), calculated from the standard deviation over trials divided by the mean response time, on both interval timing tasks (3 seconds – Control: 35.33% ± 3.50% vs. PD: 52.84% ± 2.85%, *p* = 0.0004; 7 seconds – Control: 14.94% ± 1.04% vs. PD: 23.32% ± 1.50%, *p* = 0.0002; Fig. 1D). CV was significantly correlated with cognitive function as measured by the Montreal Cognitive Assessment (MOCA) for both the 3-second (Pearson’s rho = -0.25, *p* = 0.017; Fig. 1E) and 7-second (Pearson’s rho = -0.53, *p* = 4 × 10^-8^; Fig. 1F) intervals. Furthermore, CV also predicted executive dysfunction as measured by the NIH Toolbox Dimension Change Card Sort/Flanker test for both the 3-second (Pearson’s rho = -0.29, *p* = 0.006; Fig. 1G) and 7-second (Pearson’s rho = -0.48, *p* = 2 × 10^-6^; Fig. 1F) intervals. Taken together, these data demonstrate that human PD patients had increased timing variability.

**Figure 1:**
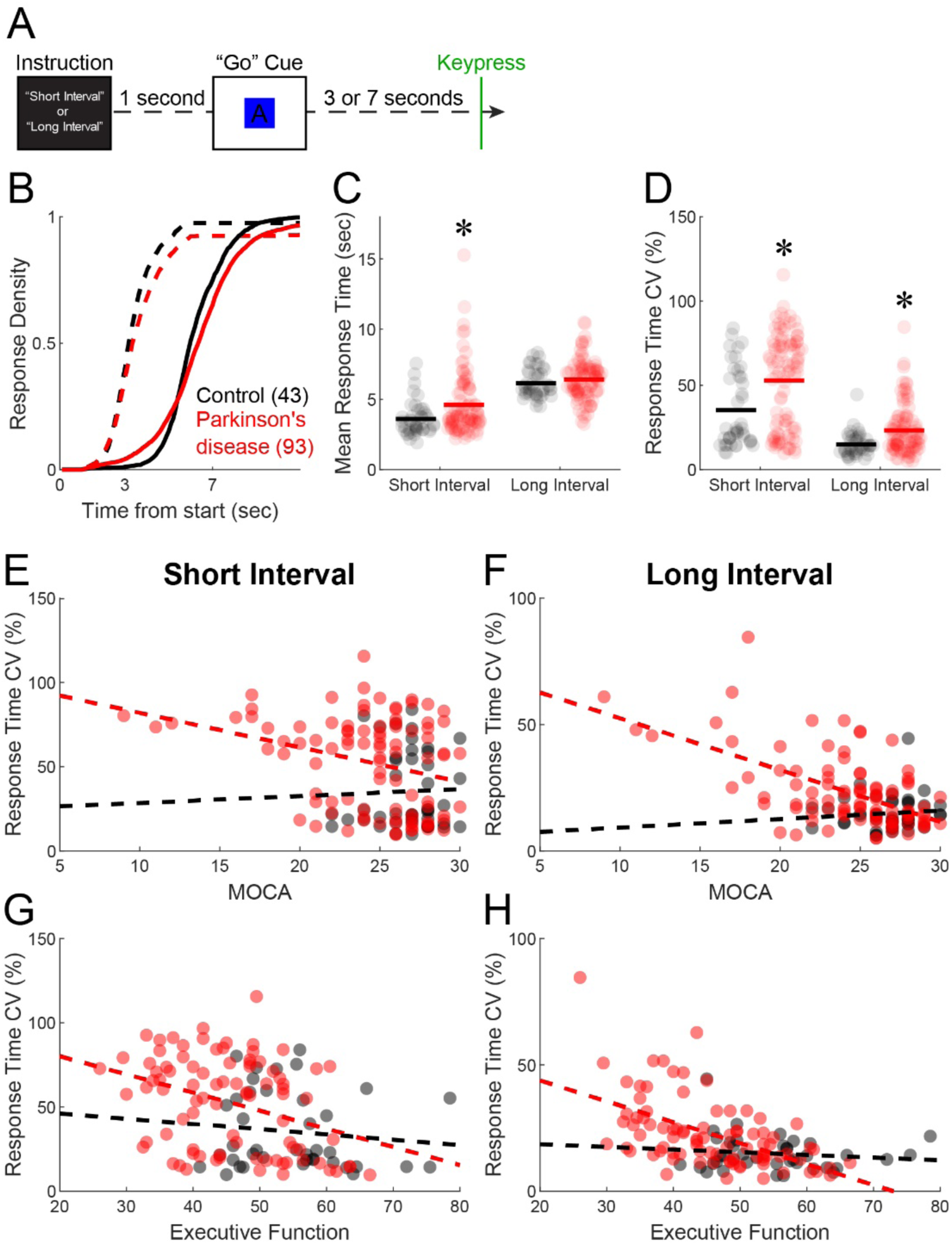
Human PD patients had increased timing variability. **A)** An interval timing task that required participants to press the spacebar when they estimated the target interval to have elapsed. **B)** Cumulative distribution of response times for 3-second (dashed) and 7-second (solid) intervals for control (black) vs. Parkinson’s disease (red). **C)** Mean response time and **D)** coefficient of variance (standard deviation/mean) of response times across participants. Response time CV for **E)** 3-second and **F)** 7-second intervals plotted vs. Montreal Cognitive Assessment (MOCA). Response time CV for **G)** 3-second and **H)** 7-second intervals plotted vs. NIH Toolbox Executive Function scores. Data from 94 PD patients and 43 controls. * *p* < 0.05.

We investigated temporal variability using a mouse-optimized interval timing task that requires mice to make a decision to switch between two nosepokes after approximately 6 seconds has passed (Fig. 2A)^18–25,35,36^. We focused on the *switch response time*, defined as the moment mice exit the first nosepoke before responding at the second nosepoke. Switch responses are a time-based decision guided by temporal control of action as no external cue indicates when to switch from one nosepoke to the other.

**Figure 2:**
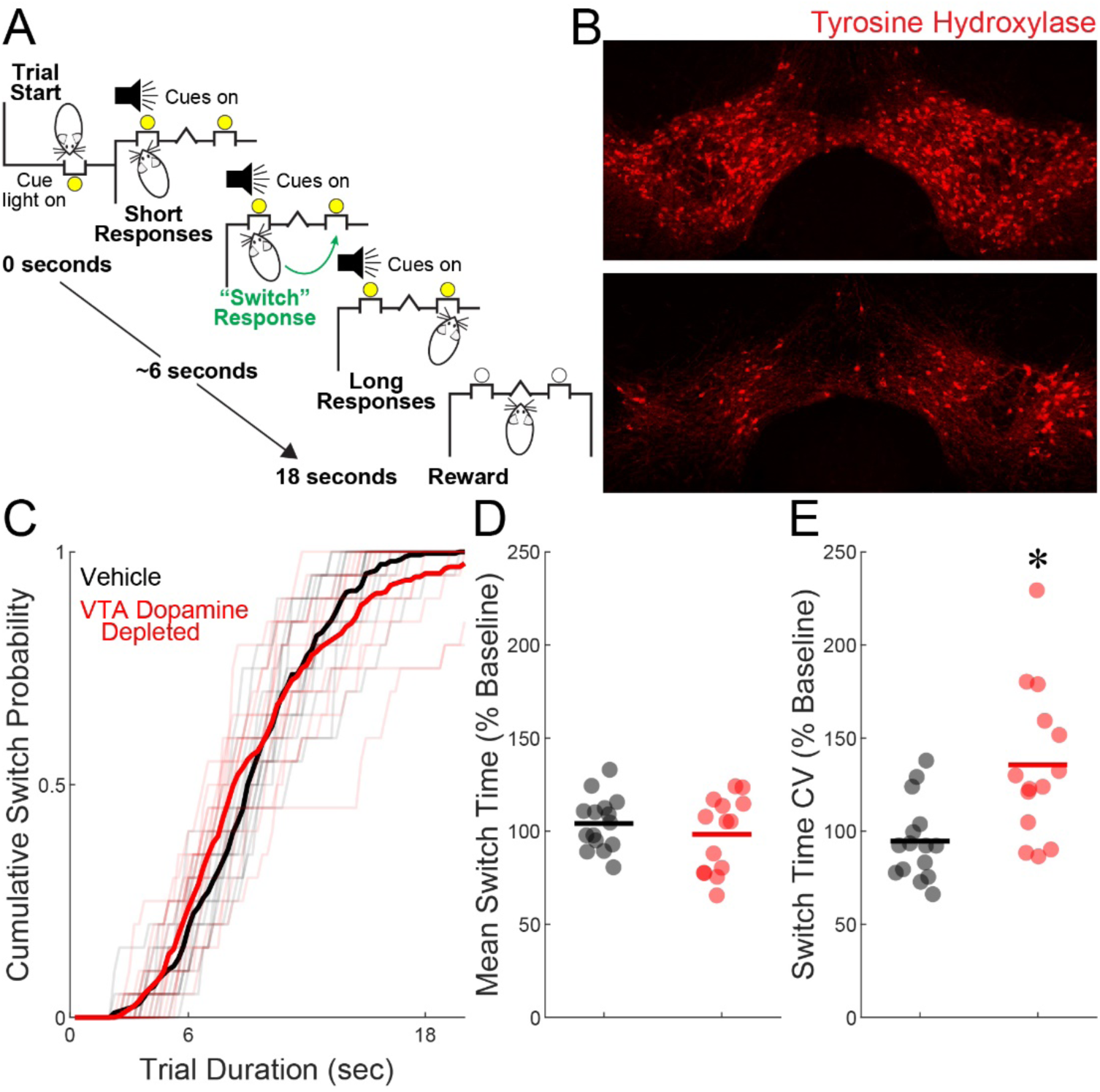
Ventral tegmental area dopamine depletion increases timing variability. **A)** Mouse-optimized interval timing in which trials are initiated at the back nosepoke. Identical cues are delivered for both randomly delivered short and long trials. On short trials, mice are rewarded for the first response after 6 seconds at the designated short nosepoke (left or right). On long trials, mice start by responding at the designated short nosepoke. When there is no reward after 6 seconds, mice switch to the designated long nosepoke (contralateral to designated short nosepoke) and wait until after 18 seconds for reward delivery. This time to switch from the short to long nosepoke is a time-based decision as in other interval timing tasks. Switch time is defined as the time of last response at the short nosepoke before responses start at the long nosepoke, and only switch trials are analyzed. **B)** Representative histological images of TH fluorescence (red) from a mouse injected with ascorbic acid vehicle (top) or 6-hydroxydopamine (6-OHDA; bottom) in the VTA. **C)** Cumulative distribution of switch responses for vehicle (black) and 6-OHDA (red) mice. **D)** Mean switch time and **E)** switch time CV, normalized to behavior prior to the lesion. Data from 15 vehicle and 14 6-OHDA mice. * *p* < 0.05.

We examined the effects of VTA dopamine depletion (Fig. 2B) on behavior in the interval timing switch task, which we have previously shown impairs interval timing^37,38^. We normalized post-surgical behavior in the switch task to pre-surgery behavior to determine magnitude change in behavior as a results of VTA dopamine depletion, similar to our previous work^37^. Data from some mice were presented in a previous report^37^ but was re-analyzed with additional mice using different methods here. The cumulative distribution of a switch for each mouse is shown in Fig. 2C. In VTA 6-OHDA treated mice, we found no significant change in mean switch time (Vehicle: 104.20% ± 3.68% (relative to presurgical baseline) vs. 6-OHDA: 98.28% ± 5.36%; *p* = 0.347; Fig. 2D), but increased timing variability as measured by the coefficient of variance (CV; Vehicle: 94.66% ± 5.50% vs. 6-OHDA: 135.69% ± 5.36%; *p* = 0.001; Fig. 2E).

There were no changes in trial initiation time (Vehicle: 58.68 ± 6.69 seconds vs. 6-OHDA: 52.38 ± 5.08 seconds; *p* = 0.449), percent of rewarded trials (Vehicle: 67.92% ± 2.26% vs. 6-OHDA: 66.95% ± 3.04%; *p* = 0.790), short responses/trial (Vehicle: 1.55 ± 0.10 vs. 6-OHDA: 1.46 ± 0.07; *p* = 0.420), long responses/trial (Vehicle: 4.50 ± 0.55 vs. 6-OHDA: 3.74 ± 0.25; *p* = 0.218), short response duration (Vehicle: 0.16 ± 0.01 seconds vs. 6-OHDA: 0.17 ± 0.01 seconds; *p* = 0.582), or long response duration (Vehicle: 0.16 ± 0.01 seconds vs. 6-OHDA: 0.17 ± 0.01 seconds; *p* = 0.367). Together, these data model deficits in human PD patients and establish a mouse model of cognitive symptoms of PD.

Next, we examined the activity of dopamine neurons using the calcium sensor GCamP6s, which was expressed in ventral tegmental dopamine neurons using DAT^IRES^-Cre mice (Fig. 3A-B). During interval timing, we found that VTA dopamine neuron GCaMP6s ΔF/F_0_ on average was modulated at trial start, drifted down over the interval, and then increased at trial end in 8 DAT^IRES^-Cre mice. We were interested in which aspects of dopamine neuron activity predicted interval timing behavior, so we analyzed GCaMP6s ΔF/F_0_ across 3 sessions from 8 DAT^IRES^-Cre mice. The mean switch time and switch time CV, separated by mouse and session, is shown in Fig 3C-D, respectively. We quantified average changes in VTA dopamine neuron GCaMP6s ΔF/F_0_ from both -0.5–0.5 seconds around trial start (Fig. 3E) and from 0.5 seconds to average switch time (Fig. 3F). We found that trial start ΔF/F_0_ strongly predicted interval timing variability (Pearson’s rho = -0.65, *p* = 0.001; Fig. 3G) and mean switch time (Pearson’s rho = -0.61, *p* = 0.002; Fig. 3H). We also found that average ΔF/F_0_ slope to switch significantly correlated with interval timing variability (Pearson’s rho = 0.42, *p* = 0.045; Fig. 3I) but did not correlate with mean switch time (Pearson’s rho = -0.21, *p* = 0.345; Fig. 3J). These data suggest that VTA dopamine neuron activity is required for and predictive of timing variability.

**Figure 3:**
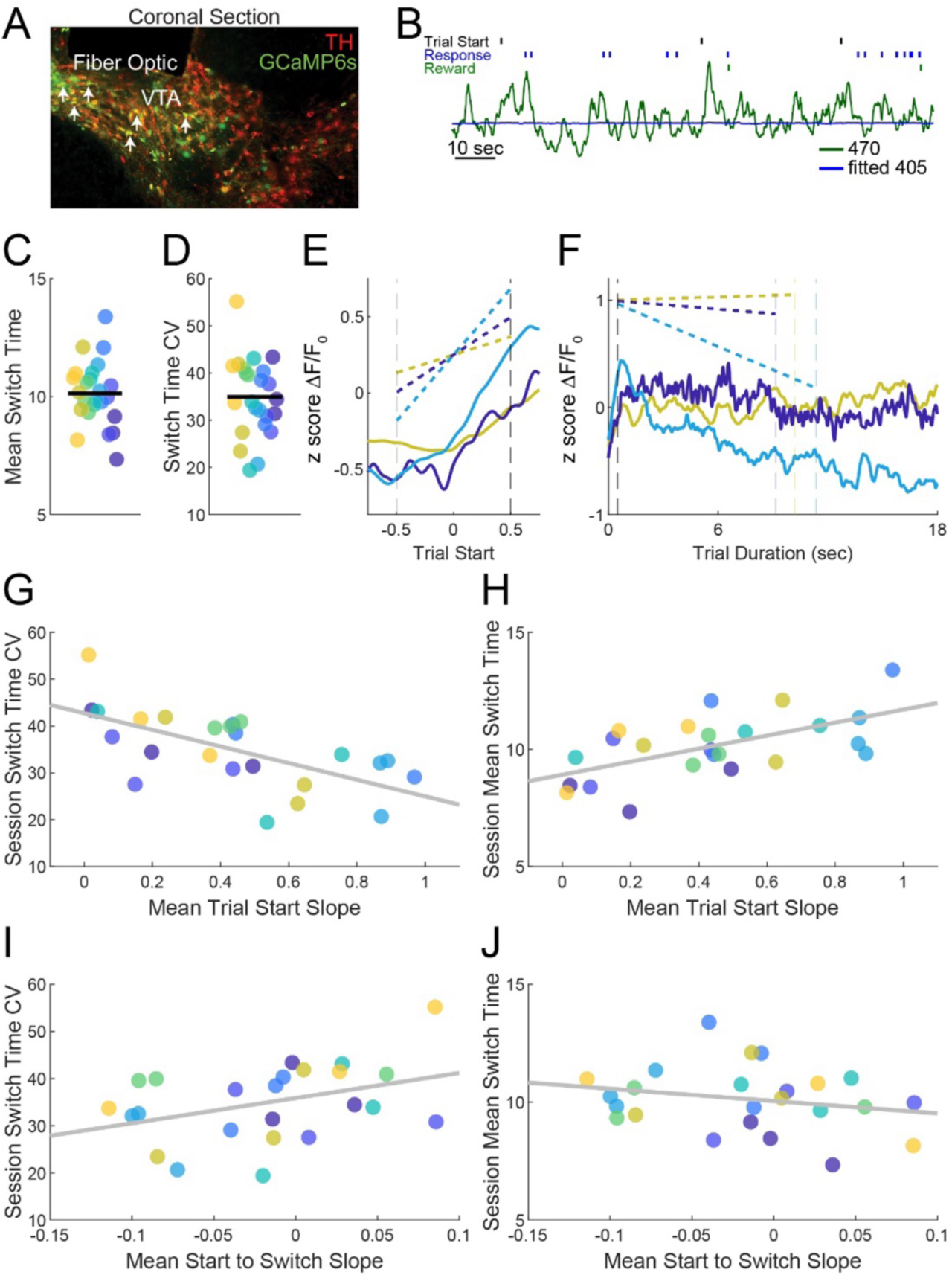
Ventral tegmental area (VTA) dopamine neuron trial-start activity predicts timing variability. **A)** Representative histological images of GCaMP6s fluorescence (green) in the VTA; tyrosine hydroxylase (TH) in red. **B)** Example GCaMP6s (470) and fitted isosbestic (405) trace over 120 seconds of a single interval timing session. **C)** Mean switch time and **D)** switch time CV from GCaMP6s ΔF/F_0_ fiber photometry sessions (each color represents a different mouse; 8 DAT^IRES^-Cre mice and 3 sessions per mouse). **E)** ΔF/F_0_ trial start slope from 3 different mice with high switch time CV (yellow), moderate switch time CV (dark blue), and low switch time CV (blue). Vertical dashed lines at -0.5 and 0.5 seconds represent the boundaries used to calculate trial start slope. The yellow, dark blue, and blue dashed lines represent the mean slope across all trials for the respective interval timing session. **F)** ΔF/F_0_ from trial start to switch from the same sessions as shown in **E**. Vertical dashed lines are at 0.5 seconds and session mean switch time. The yellow, dark blue, and blue dashed lines represent the mean start to switch slope across all trials for the respective interval timing session. **G)** Trial start slope was inversely correlated with CV and **H)** correlated with mean switch time. **I)** There was also a correlation between start-to-switch slope and CV, but **J)** start-to-switch slope did not correlate with mean switch time. Data from 3 sessions in 8 DAT^IRES^-Cre mice. * *p* < 0.05.

We then tested the hypothesis that stimulating VTA dopamine neurons would decrease timing variability. We expressed channelrhodopsin (ChR2) in the ventral tegmental area of DAT^IRES^-Cre mice. In non-dopamine depleted mice (Fig. 4A), we found that stimulating VTA dopamine neurons at 4-Hz for the 2 seconds, which corresponds to 8 laser pulses, after trial start improved interval timing by decreasing variability (No Stimulation: 36.47% ± 1.17% vs. 4-Hz Stimulation 31.54% ± 1.07%, *p* = 0.001; Fig. 4B-C). There was no change in mean switch time (No Stimulation 9.47 ± 0.22 seconds vs. 4-Hz Stimulation 9.65 ± 0.18 seconds, *p* = 0.439; Fig. 4D). Interestingly, 20-Hz stimulation for the same 2 seconds after trial start did not change timing variability (No Stimulation 37.21% ± 1.86% vs. 20-Hz Stimulation 37.70% ± 0.73%, *p* = 0.794) or mean switch time (No Stimulation 8.74 ± 0.25 seconds vs. 20-Hz Stimulation 8.98 ± 0.29 seconds, *p* = 0.518).

**Figure 4:**
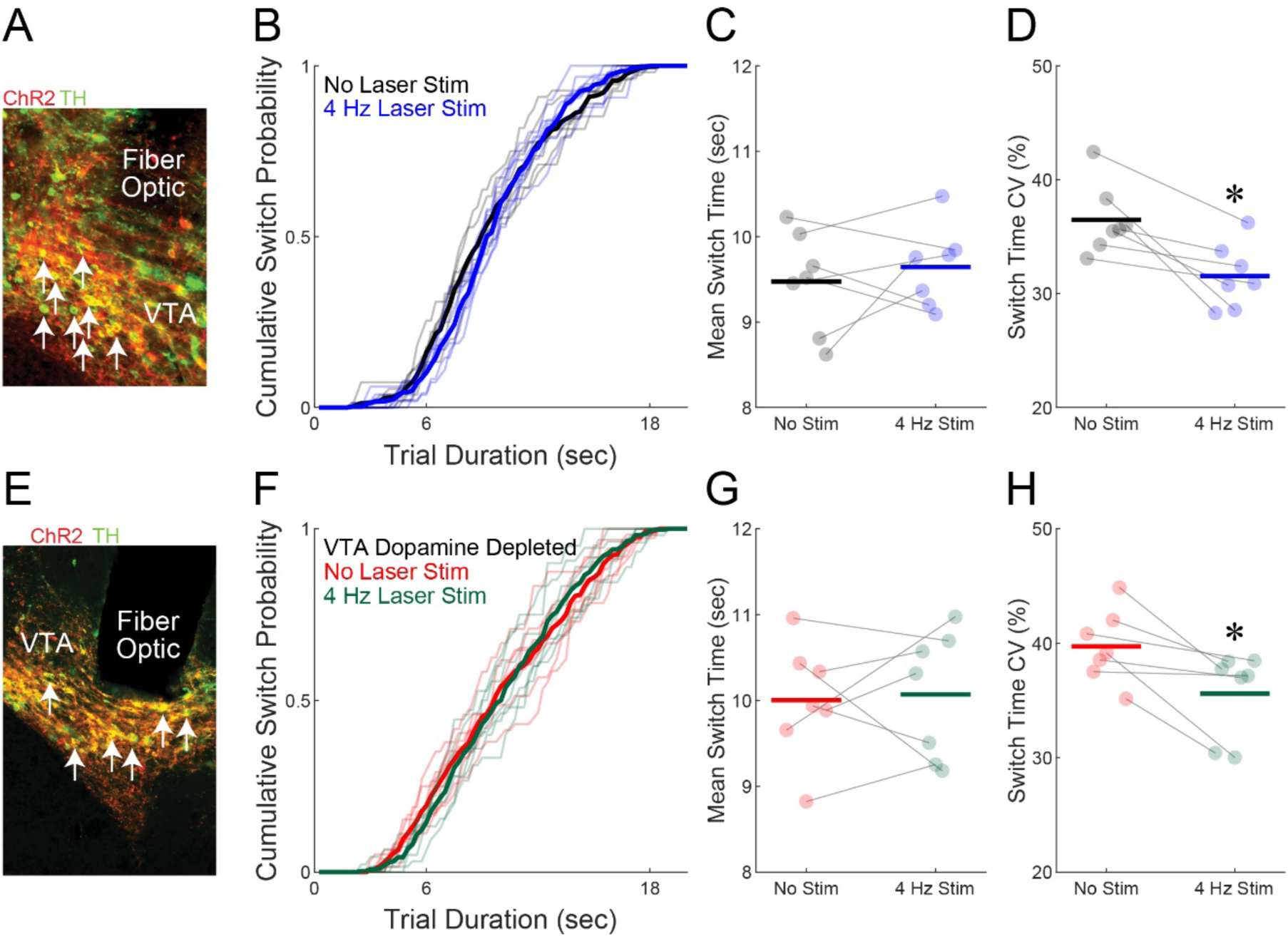
Stimulating ventral tegmental area (VTA) dopamine neurons improves timing variability. **A)** Representative histological images of fiber optics and channelrhodopsin (ChR2; red) in the VTA; tyrosine hydroxylase (TH) in green. **B)** Cumulative switch responses for no stimulation (black) and 4-Hz VTA dopamine neuron stimulation (blue). **C)** There was no change in mean switch time but **D)** there were reliable improvements in timing by decreasing interval timing CV. **E)** In dopamine-depleted mice, representative histological images of fiber optics and ChR2 (red) in the VTA; TH in green. **F)** Cumulative switch responses for no stimulation (red) and 4-Hz VTA dopamine neuron stimulation (green). **G)** There was no change in mean switch time, but **H)** there were reliable improvements in timing by decreasing interval timing CV. Data from 14 DAT^IRES^-Cre mice (7/experiment). * *p* < 0.05.

No effects were observed in control animals with identical stimulation parameters expressing a virus without opsins. There was no change in timing variability at either 4-Hz (No Stimulation 34.03 ± 2.30 vs. 4-Hz Stimulation 34.67 ± 2.33, *p* = 0.702) or 20-Hz (No Stimulation 31.57 ± 2.09 vs. 20 Hz Stimulation 33.77 ± 1.28, *p* = 0.287). There was no change in mean switch time at either 4-Hz (No Stimulation 10.21 ± 0.29 vs. 4-Hz Stimulation 10.29 ± 0.41, *p* = 0.760) or 20-Hz (No Stimulation 9.88 ± 0.11 vs. 20-Hz Stimulation 9.84 ± 0.27, *p* = 0.834).

We were interested if VTA dopamine neuron stimulation could compensate for cognitive symptoms of PD. We turned to mice with intermediate dopamine depletion, in which some dopamine neurons remain. Notably, even in human Parkinson’s disease, ∼30% of dopamine neurons are spared at the end of their disease^39^. Accordingly, we expressed ChR2 in the VTA of DAT^IRES^-Cre mice with dopamine depletion (Fig. 4E). Similar to non-dopamine depleted mice, we found to our surprise that VTA dopamine neuron stimulation at 4-Hz improved interval timing by decreasing timing variability (No Stimulation 39.72% ± 1.20% vs. 4-Hz Stimulation 35.60% ± 1.41%, *p* = 0.003; Fig. 4F-G) but did not change mean switch time (No Stimulation 10.00 ± 0.26 seconds vs. 4-Hz Stimulation 10.07 ± 0.28 seconds, *p* = 0.816; Fig. 4H). There was no effect of 20-Hz stimulation on timing variability (No Stimulation 41.97% ± 1.64% vs. 20-Hz Stimulation 43.58% ± 1.81%, p = 0.304) or mean switch time (No Stimulation 9.43 ± 0.33 seconds vs. 20-Hz Stimulation 9.21 ± 0.30 seconds, p = 0.486). Together, these data demonstrate that VTA dopamine neurons control timing variability.

## DISCUSSION

We studied behavioral variability using interval timing. We report three main results. First, PD patients have increased timing variability that predicts cognitive impairment and executive function. Second, we can model this increased timing variability in mice with ventral tegmental area dopamine depletion. Third, VTA dopamine neuron calcium dynamics at the start of temporal intervals is highly predictive of temporal variability across animals. Finally, we find that stimulating VTA dopamine neurons improved interval timing by decreasing temporal variability in intact and dopamine-depleted mice. These data establish that targeting VTA dopamine in mice may model aspects of cognitive dysfunction in human PD^6^ and elucidate neuronal circuits controlling behavioral variability.

VTA neurons play a key role in motivated behavior. VTA dopamine neurons are heterogenous, projecting to the prefrontal cortex and nucleus accumbens along with other brain stem and subcortical nuclei. Of these, only prefrontal networks are reliably involved in executive dysfunction^40–42^. While much focus has been placed on the role of VTA in motivation and reward^43,44^, here we demonstrate a role of VTA dopamine in behavioral variability, independent of reward or motivational effects. Our work is consistent with a role for dopamine in executive function^45^. Indeed, depleting prefrontal dopamine profoundly impairs executive functions such as working memory and interval timing^20,46,47^, which requires working memory for temporal rules and attention to the passage of time. Of note, VTA dopaminergic circuits are complex, as VTA dopamine neurons can co-release GABA and glutamate^48,49^, and have distinct transcriptional markers such as *Calb1*^50,51^. Future work will need to map this complexity, but the present work is clinically significant because VTA neurons degenerate in PD^39^ and may contribute to cognitive symptoms in PD^52^.

We found that 4-Hz VTA dopamine neuron stimulation improved interval timing by decreasing variability. This finding is supported by similar work showing that low-frequency prefrontal stimulation of neurons expressing D1-type dopamine receptors improves interval timing^33,34,47^ and that 4-Hz cerebellar stimulation improves interval timing in rodent models^53^. In humans, we found that 4-Hz subthalamic nucleus stimulation improved interval timing^54^. Of note, trial start-evoked 4-Hz activity is linked with dopamine, deficits in interval timing, and cognitive symptoms of PD^17,17,33,55,56^. Work by our group and others have found that ∼4-Hz deep-brain stimulation of the subthalamic nucleus can improve executive function in PD^54,57–59^. It is unclear how 4-Hz subthalamic nucleus stimulation affects ventral tegmental area or cortical circuits. Importantly, our stimulation was triggered by the start of the interval, while other stimulation paradigms are continuous. Future work will resolve how low-frequency stimulation affects cortical networks, which could be critical to advancing new therapies.

Our work has several limitations. First, we did not explore VTA dopaminergic neuron subtypes or projections^49,50^. Second, we did not explore which cortical or subcortical circuits and neurons are responsible for changing variability, although such effort would require a broad coordinated approach. Finally, we did not deliver brain stimulation to the ventral tegmental area in humans, which would require new surgical planning and dedicated clinical trials. Despite these limitations, our work advances a new role of VTA dopamine neurons in cognitive variability, which is distinct from its traditional role in motivation and reward. Our work will inspire new markers and therapies for human diseases that degrade cognitive variability that may lead to cognitive impairment and dementia.

## Data and Code Availability

All code and data will be available upon final publication at: http://narayanan.lab.uiowa.edu/article/datasets.

## Author Contributions

MAW and NSN designed the animal experiments. AS, RCC, AIE, and NSN designed the human experiments. MAW, KS, EET, MMC, and MO performed all animal experiments. MAW, KS, and EET independently verified histological targeting and performed histological immunofluorescent analysis. MAW, ASB, YK, and NSN performed all statistical analyses. MAW and NSN wrote the manuscript, and all authors reviewed and revised the manuscript.

